# SARS-CoV-2 exposure in Norway rats (*Rattus norvegicus*) from New York City

**DOI:** 10.1101/2022.11.18.517156

**Authors:** Yang Wang, Julianna Lenoch, Dennis Kohler, Thomas J. DeLiberto, Cynthia Tang, Tao Li, Yizhi Jane Tao, Minhui Guan, Susan Compton, Caroline Zeiss, Jun Hang, Xiu-Feng Wan

**Author notes:** Xiu-Feng Wan; Thomas J. DeLiberto, **Email:**. **Author Contributions:** XFW, JL, DK, and TD conceived the experiments. XFW, JL, DK, TD, and YW designed the experiments. JL, DK, and TD contributed to the wild rat capture. YW, TL, JT, and MG performed the experiments. SC and CZ provided study materials. YW, CT, JT, TL, and JH analyzed the data. YW and XFW prepared the original draft. All authors did substantial contribution to revision of this paper.

## Abstract

Millions of Norway rats (*Rattus norvegicus*)inhabit New York City (NYC), presenting the potential for transmission of SARS-CoV-2 from humans to rats and other wildlife. We evaluated SARS-CoV-2 exposure among 79 rats captured from NYC during the fall of 2021. Results showed that 13 of 79 rats (16.5%) tested IgG or IgM positive, and partial genomes of SARS-CoV-2 were recovered from four rats that were qRT-PCR positive. Using a virus challenge study, we also showed that Alpha, Delta, and Omicron variants can cause robust infections in wild-type Sprague Dawley (SD) rats, including high level replications in the upper and lower respiratory tracts and induction of both innate and adaptive immune responses. Additionally, the Delta variant resulted in the highest infectivity. In summary, our results indicated that rats are susceptible to infection with Alpha, Delta, and Omicron variants, and rats in the NYC municipal sewer systems have been exposed to SARS-CoV-2. Our findings highlight the potential risk of secondary zoonotic transmission from urban rats and the need for further monitoring of SARS-CoV-2 in those populations.

**Importance:** Since its emergence causing the COVID-19 pandemic, the host tropism expansion of SARS-CoV-2 raises a potential risk for reverse-zoonotic transmission of emerging variants into rodent species, including wild rat species. In this study, we presented both genetic and serological evidence for SARS-CoV-2 exposure in wild rat population from New York City, and these viruses are potentially linked to the viruses during the early stages of the pandemic. We also demonstrated that rats are susceptible to additional variants (i.e., Alpha, Delta, and Omicron) predominant in humans and that the susceptibility to different variants vary. Our findings highlight the potential risk of secondary zoonotic transmission from urban rats and the need for further monitoring of SARS-CoV-2 in those populations.

## Introduction

As of October 10, 2022, severe acute respiratory syndrome coronavirus 2 (SARS-CoV-2), the virus responsible for coronavirus disease 2019 (COVID-19), has caused approximately 621 million human cases and 6.6 million deaths globally (1). In addition to humans, a wide range of wild, domestic, and captive animals were documented with exposure to SARS-CoV-2, such as deer, mink, otters, ferrets, hamsters, gorillas, cats, dogs, lions, and tigers (2–4). SARS-CoV-2 in farmed mink was shown to cause infections in humans (5), highlighting mink as a potential reservoir for secondary zoonotic infections.

SARS-CoV-2 has undergone rapid evolution, and a large number of genetic variants have been identified, including several variants of concern (VOC), such as Alpha (B.1.1.7 lineage), Beta (B.1.351 lineage), Gamma (P.1 lineage), Delta (B.1.617.2 and AY sublineages) and Omicron (B.1.1.529 and BA sublineages). The Alpha, Beta, and Gamma variants were reported to have acquired substitutions at the receptor-binding domain (RBD) of the spike protein that allowed for infectivity in mice and/or rats (6–9). The tropism expansion of SARS-CoV-2 raises a potential risk for reverse-zoonotic transmission of emerging variants into rodent species, including wild mouse and rat species (10). Two independent SARS-CoV-2 serosurveillance studies among wild rats from sewage systems in Belgium (late fall of 2020) and Hong Kong (spring of 2021) suggested possible exposure of these animals to SARS-CoV-2, but no viral RNA was detected (11, 12). With new SARS-CoV-2 variants continuing to emerge, it is still unknown whether the more recent variants of concern (e.g., Delta and Omicron) are infectious to rats.

In this study, we evaluated the capability of Delta and Omicron variants to infect rats (*Rattus norvegicus*) and investigated the exposure of rats to SARS-CoV-2 in New York City (NYC), New York, United Sates.

## Results

### Detecting SARS-CoV-2 virus in NYC rats

To evaluate whether wild rats have been exposed to SARS-CoV-2, we conducted SARS-CoV-2 surveillance in Norway rats (*Rattus norvegicus*) in NYC from September 13-November 16, 2021, when the Delta variant was predominant in humans. A total of 79 rats inhabiting three sampling sites in Brooklyn, NYC were captured and sampled. Using ELISA, we identified 9 out of 79 (11.4%) IgG-positive rat serum samples and 4 IgM-positive samples (5.1%) against both Wuhan-Hu-1 spike protein and RBD (Table 1). All 13 seropositive samples were subjected to microneutralization assays against the B.1 lineage and the Alpha and Delta variants. However, all samples were negative for neutralizing antibodies. As a negative control, we used ELISA to examine 9 negative serum samples from uninfected SD rats and 6 serum samples from SD rats infected with rat coronaviruses, Sialodacryoadenitis Virus or Parker’s Rat Coronavirus (13); none exhibited IgG or IgM positivity against either spike protein or RBD (Data not shown).

**Table 1.**
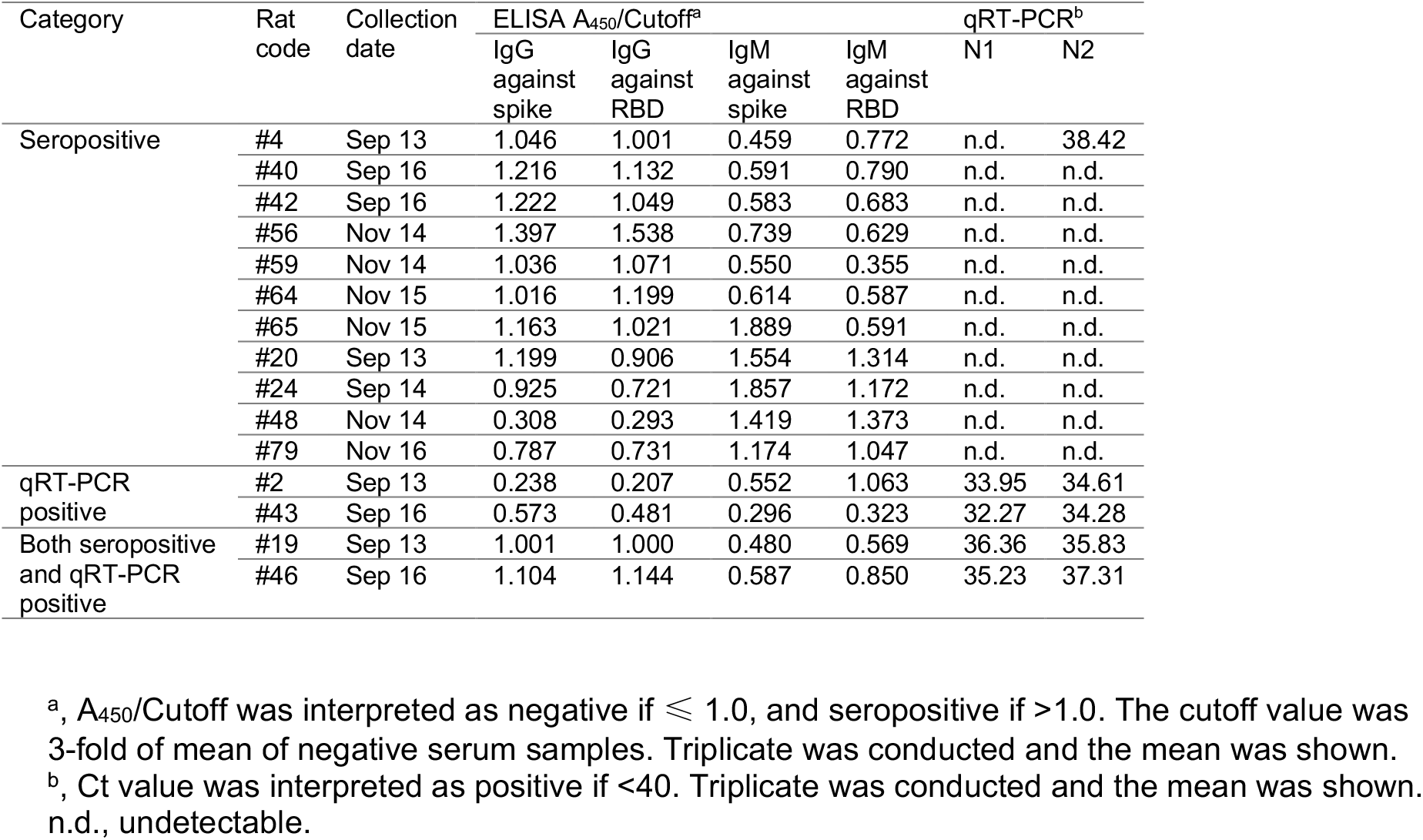
Information on rats collected in Brooklyn of NYC with conclusive seropositive or qRT-PCR positive samples

Of all the tissues analyzed from the 79 rats, only four lung samples were positive by qRT-PCR against both N1 and N2 primers using the CDC SARS-CoV-2 diagnostic panel (Table 1). The control group with RNA from 6 different strains of rat coronaviruses remained negative. It is noteworthy that two out of these four rats (Rat #2 and #19) were both seropositive and viral RNA-positive. In addition, we had seven inconclusive samples which were tested positive on either N1 or N2 primer but not both. However, viruses failed to be recovered from Vero E6, 293FT/hACE2+TMPRSS, rat lung epithelial (L2), or rat lung tracheal epithelial cell lines.

After subjecting these four qRT-PCR-positive samples to SARS-CoV-2 genome sequencing, partial SARS-CoV-2 genome was identified in all samples with a viral genome coverage of 1.6% to 21.3% (Table S1). Both molecular characterization and phylogenetic analyses on these partial genomes suggested that viruses in these rats are associated with genetic lineage B, which was predominant in NYC in the spring of 2020 during the early pandemic period (Fig. 1).

**Figure 1.**
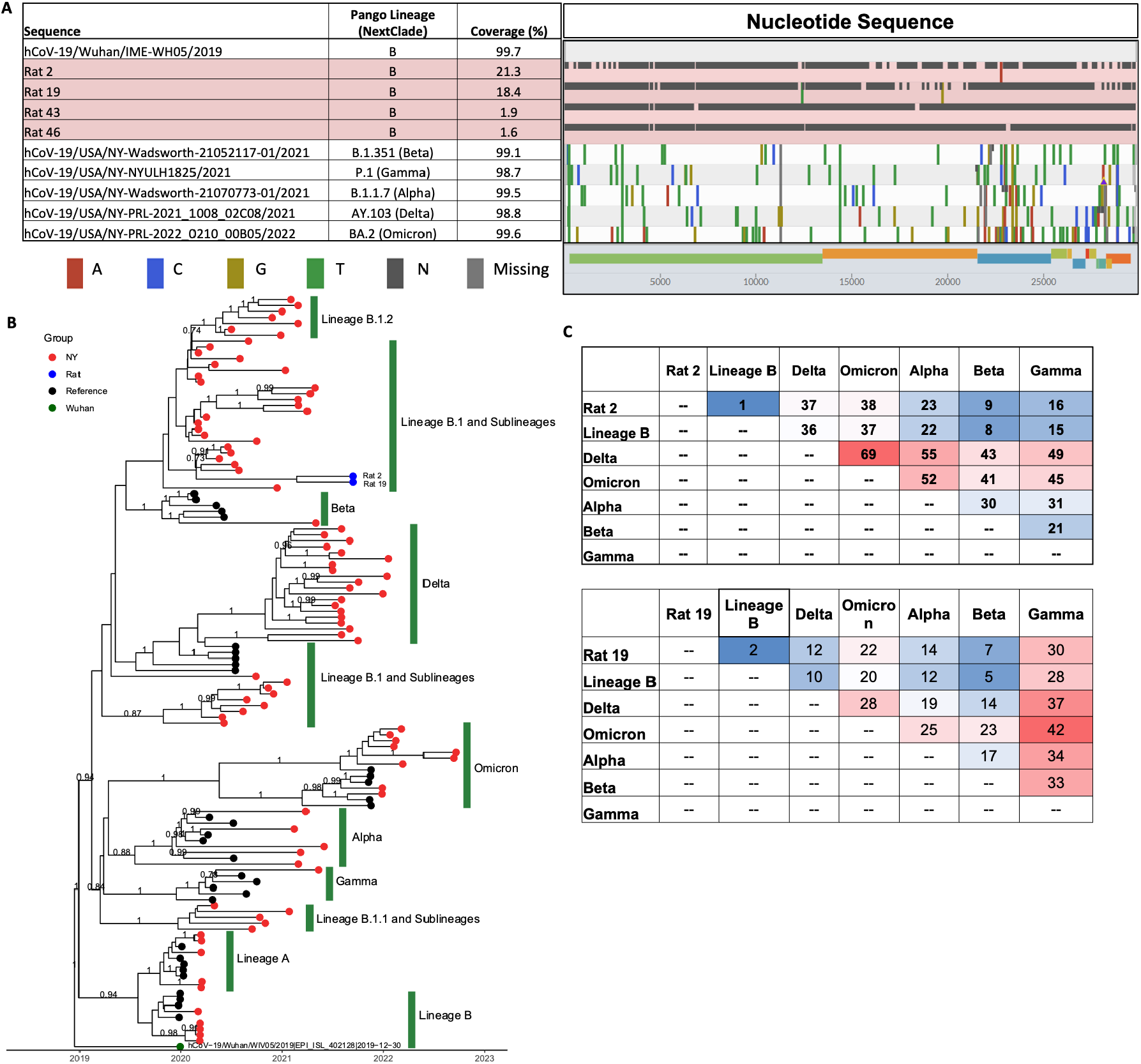
SARS-CoV-2 genomic sequencing in wild rats. (A) SARS-CoV-2 genomes found in rats in comparison with reference wild-type virus and variants of concern. Analyses were performed and visualized using https://clades.nextstrain.org. Reference sequences were downloaded from GISAID. (B) Phylogenetic tree of rat SARS-CoV-2 sequences with reference sequences from wild-type viruses and variants of concern. Branches with posterior probability ≥ 0.7 are labeled. (C) Distance matrices of regions covered by each rat-derived SARS-CoV-2 genome. Lineage B is represented by hCoV-19/Wuhan/IME-WH05/2019|EPI_ISL_529217|2019-12-30, Delta by hCoV-19/USA/NY-Wadsworth-21052117-01/2021|EPI_ISL_2278740|2021-05-01, Omicron by hCoV-19/USA/NY-NYULH1825/2021|EPI_ISL_2427410|2021-05-11, Alpha by hCoV-19/USA/NY-Wadsworth-21070773-01/2021|EPI_ISL_2868594|2021-05-31, Beta by hCoV-19/USA/NY-PRL-2021_1008_02C08/2021|EPI_ISL_5285364|2021-10-03.

In addition, we subjected these four qRT-PCR-positive and two additional inconclusive samples to pan-viral target hybridization enrichment sequencing. Presence of SARS-CoV-2 sequences were found in three out of four sequenced qRT-PCR-positive samples (Rats# 2, 19, and 43) and one of two inconclusive samples (Rat# 38). No sequence data was obtained for the qRT-PCR-positive sample from Rat# 46. Of interest, rat coronavirus was detected in another inconclusive sample (Rat# 30) (Table S2). The identified SARS-CoV-2 or rat coronavirus reads aligned with a number of genes across the genomes.

### Rats displayed varying susceptibility to SARS-CoV-2 variants

The Alpha variant emerged in late 2020 and quickly became a dominant SARS-CoV-2 variant in NYC; subsequently, the Delta and Omicron variants predominated in NYC starting in June 2021 and December 2021, respectively (Fig. 1A). To investigate whether these SARS-CoV-2 variants are capable of infecting rats, we intranasally challenged 6-week-old wild-type SD rats with Alpha, Delta, or Omicron variants and collected tissues at 2- and 4-days post-infection (dpi) (Fig. 2B). Compared to the Wuhan-Hu-1 strain, the Omicron variant used in the challenge study possesses the same N501Y substitution as the Alpha variant and 16 additional substitutions, whereas the Delta variant does not possess N501Y, but contains the L452R and T478K substitutions (Fig. 2C).

**Figure 2.**
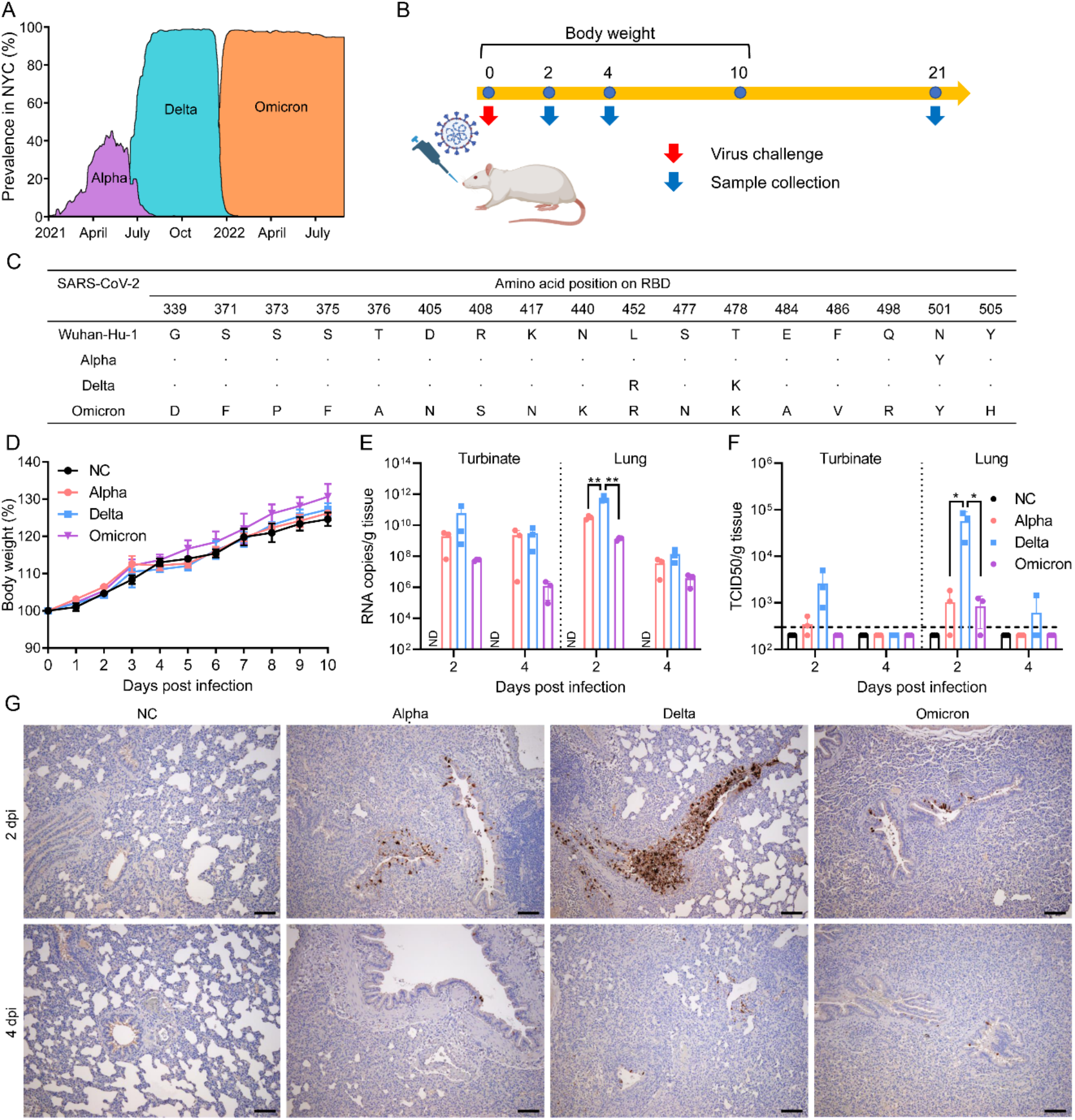
SD rats are susceptible to infection of Alpha, Delta, and Omicron variants. (A) The prevalence of Alpha, Delta, and Omicron variants in NYC. The figure was adapted from https://outbreak.info. (B) Scheme of the virus challenge experiment using 6-week-old SD rats. (C) Amino acid changes of Alpha, Delta, and Omicron variants across RBD compared to Wuhan-Hu-1 (NCBI access No.: MN908947.3). (D) Body weight of rats mock infected or infected with either Alpha, Delta, or Omicron variant, Viral RNA copies (E) and infectious viral titers (F) in the turbinate and lungs from rats infected with either Alpha, Delta, or Omicron variant at 2 or 4 dpi. *, *p* < 0.05; **, *p* < 0.01. (G) Detection of SARS-CoV-2 nucleocapsid protein at bronchial epithelial cells by immunohistochemistry at 2 and 4 dpi. Scale bar, 100 μm.

At 2 and 4 dpi, high levels of viral RNA were detected in both turbinate and lungs, and infectious viral titers were detected in turbinate and/or lungs, although no body weight loss or other clinical signs were observed in the rats with any of the variants (Fig. 2D-F). In particular, the lungs from the rats infected with the Delta variant showed both the highest RNA copies and the highest infectious viral titers at 2 dpi (RNA copies: *p*=0.0081 and 0.0060 for Delta vs. Alpha and Delta vs. Omicron, respectively; infectious viral titers: *p*=0.0287 and 0.0283 for Delta vs. Alpha and Delta vs. Omicron, respectively). In addition, antigen expression was detected in the lungs of all rats infected with any variant at 2 or 4 dpi (Fig. 2F). In line with the viral titers, the rats infected with the Delta variant showed the highest antigen expression in the lungs compared to those infected with other variants (Fig. 2G).

To assess the innate and adaptive immune response induced by the virus infection in rats, we determined the cytokine/chemokine expression in the lungs at 2 and 4 dpi and the antibody titers at 21 dpi. The results showed that all infections induced pro-inflammatory cytokine/chemokine expression (i.e., IFN-β, IFN-γ, TNF-α, IL-1α, IL-1β, IL-6, CCL-2, IP-10, IL-10) particularly at 2 dpi (Fig. 3A). The expression of all the cytokines/chemokines induced by the Delta variant was higher than those induced by Alpha and/or Omicron variants.

**Figure 3.**
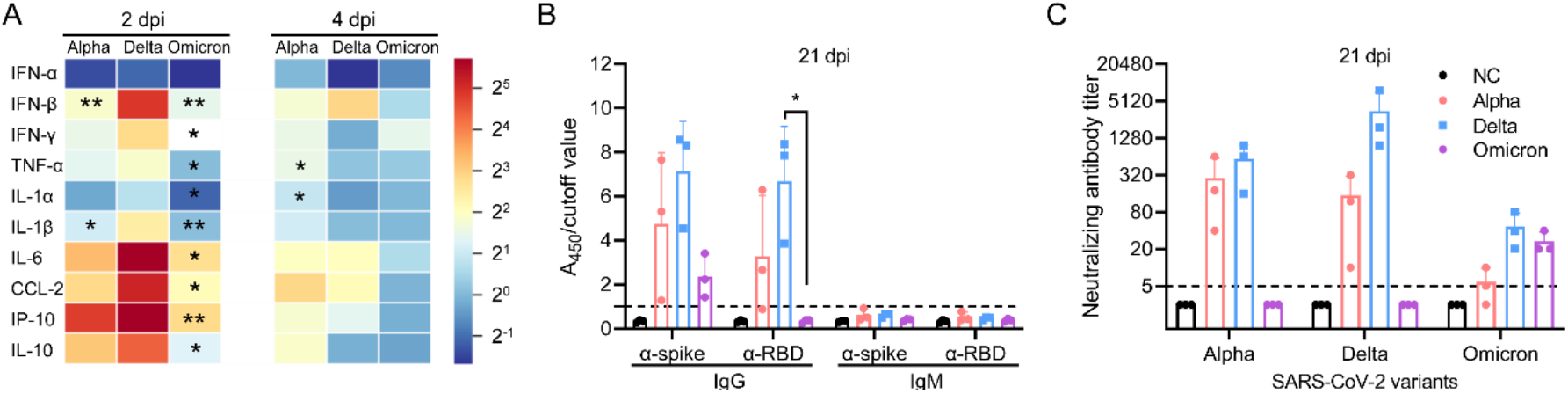
Innate and adapted immune responses induced by SARS-CoV-2 infection in SD rats. (A) Pro-inflammatory cytokine/chemokine expression in lungs from rats infected with either Alpha, Delta, or Omicron variant. Asterisks indicate the significant difference between the indicated variant and Delta. *, *p* < 0.05; **, *p* < 0.01. (B) Wuhan-Hu-1 spike protein or RBD specific IgG or IgM antibody titers. A_450_/Cutoff was interpreted as negative if ≤ 1.0, and seropositive if >1.0. The cutoff value was 3-fold of mean of negative serum samples. (C) Alpha, Delta, or Omicron specific neutralizing antibody titers induced by mock-infected rats or rats infected with either Alpha, Delta, or Omicron variant.

Regarding the adaptive immune response, both IgG antibodies and neutralizing antibodies were detected for all three variants at 21 dpi; however, IgM antibodies were not detected in any rats regardless of the variant used (Figures 3B and 3C). There was no significant difference between Alpha and Delta variants in the IgG antibody titers against Wuhan-Hu-1 spike protein or RBD. However, Delta showed significantly higher anti-RBD IgG titers than Omicron. The homologous neutralizing antibody titers induced by the Delta variant were significantly higher than those induced by Alpha or Omicron (p=0.0441 and 0.0040, respectively). These results indicated that all the three variants can infect SD rats and induce innate and adaptive immune responses, and among these three variants, the Delta variant replicates more efficiently than the Alpha and Omicron variants in rats.

To detect potential host-adapted mutations, we sequenced the lung tissues from the rats challenged with Alpha, Delta, and Omicron. Results suggested there were no adapted amino acid substitutions along the RBDs across the three testing variants. However, N74K (N-terminal domain) on the spike protein was observed in all animals challenged by Alpha, and P681R (SD1/2) and D950N (heptapeptide repeat sequence 1) of spike in all animals challenged by Delta (Table S2). In addition, additional amino acid substitutions in non-structural proteins NSP6, NSP13, and nucleoprotein were observed in some animals challenged by Alpha or Delta. Of interest, no adapted mutations were observed in the animals challenged by Omicron.

### Structural modeling between RBD of SARS-CoV-2 variants and rat, mouse, and human ACE2

To explain the relative replication efficiency of the three SARS-CoV-2 viruses in SD rats, we computationally modeled the interaction between rat ACE2 and RBD from Alpha, Delta and Omicron variants (Fig. 4), as virus-receptor interaction is often an important virulence determinant. In our structural models, residue 452 does not directly engage with rat ACE2, but it is surrounded by a large number of residues nearby (Fig. 4C). Therefore, the L452R mutation in the Delta variant could alter the structure conformation of the adjacent β-strand at the ACE2 interface and thus indirectly modulate ACE2 binding affinity (Fig. 4C). Indeed, in vitro binding assays indicated that the RBD of the Delta variant, which has L452R/T478K double mutations, binds rat ACE2 with a >2-fold stronger affinity than RBD of the prototype virus (14). The enhanced binding of the Delta RBD to rat ACE2 is likely due to L452 alone, because residue 478 is distant from other amino acids, and T478K was found to have no significant effect on binding to mouse ACE2, which is a close homolog of rat ACE2 (15). The Alpha variant also replicates well in rats but is slightly less efficient than Delta. Our structure model shows that the single mutation N501Y in Alpha RBD makes a favorable interaction with H353 in the rat ACE2, with the aromatic side chain of Y501 stacked against the side chain of H453 (Fig. 4D). In vitro binding assays confirmed that the Alpha RBD binds rat ACE2 with a >2-fold stronger affinity than RBD of the prototype virus (14), consistent with our structural analysis.

**Figure 4.**
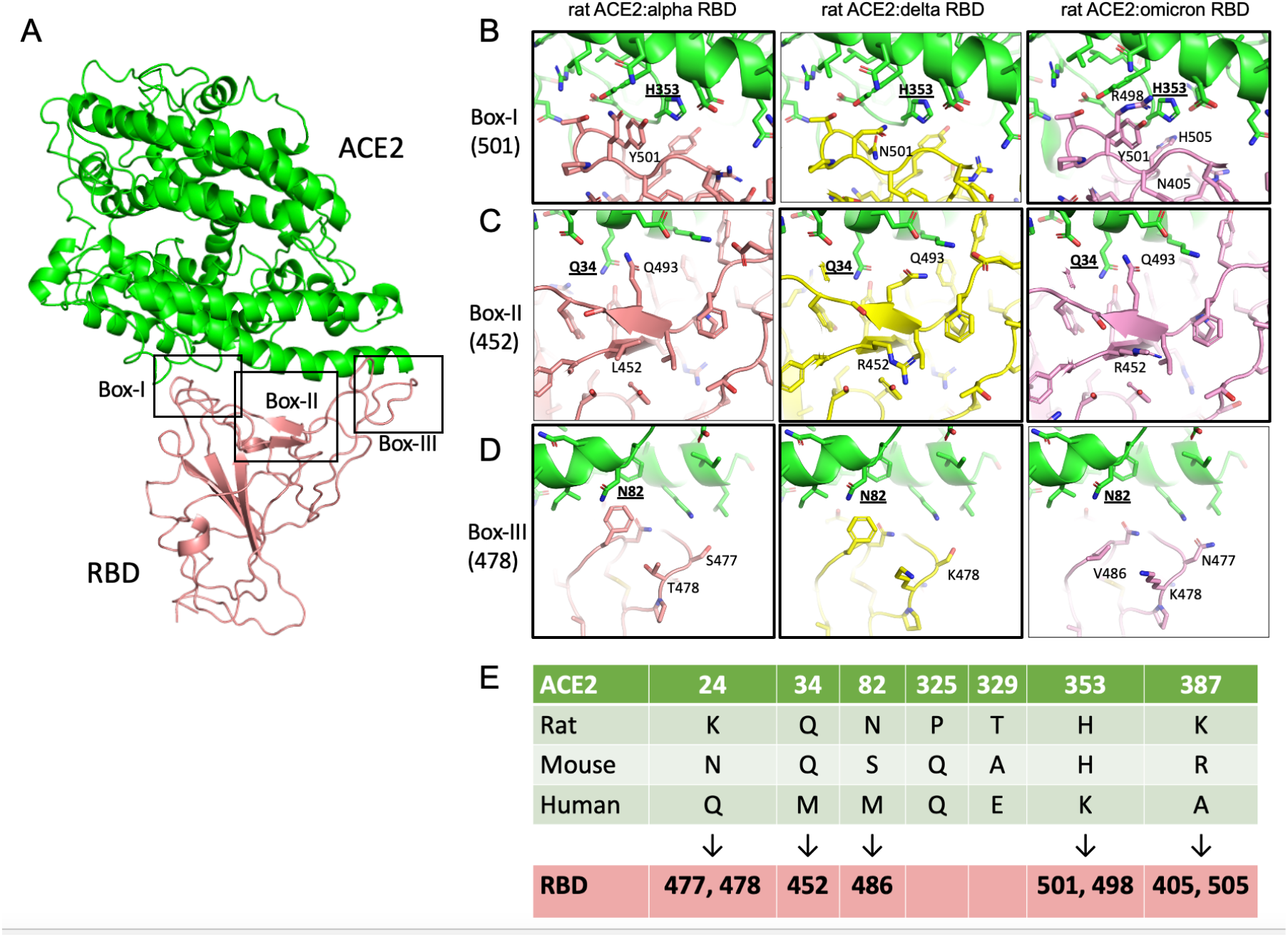
Interactions between the receptor binding domain (RBD) of SARS-CoV-2 variants Alpha, Delta, and Omicron and the rat ACE2. (A) Rat ACE2 in complex with RBD. The three major contact sites in box-1, box-2 and box-3 are shown in subpanels B, C, and D, respectively. Interactions mediated by alpha, delta and omicron RBDs are compared side-by-side. Black thick outlines highlighted favorable interactions. (E) A list of ACE2 amino acid variations between rat, mouse and human at the RBD interface. Many RBD mutations in alpha, delta and omicron variants are located near host-specfic ACE2 residues, as indicated by black arrows.

The Omicron variant has many mutations in its RBD compared to the prototype virus (Fig. 2C). Among these mutations, eight are located near the ACE2 binding interface, including residues 405, 452, 477, 478, 486, 498, 501, and 505. Close inspection of these residues shows favorable interactions by residues R452, N477, R498, Y501, and H505 compared to their corresponding ones in the prototype strain. Residues D405 and K478 are somewhat distant from ACE2, while V486 appears to weaken the interaction with rat ACE2 compared to F486 in other SARS-CoV-2 viruses (Fig. 3D).

Taken together, the Alpha, Delta and Omicron variants seem to have enhanced binding to the rat ACE2 compared to the prototype virus.

## Discussion

Both serological and molecular data from this study suggested the rats from NYC were exposed to SARS-CoV-2. We found that of the tested rats, 16.5% were seropositive and 5.1% were qRT-PCR positive to SARS-CoV-2, which showed a higher exposure frequency than previous reports (11, 12). Genomic analyses suggested that the viruses in the rats that we collected were associated with the B lineage virus. We speculate SARS-CoV-2 exposure could have occurred during the early stages of the pandemic when the B lineage virus was predominant in NYC. This is supported by a recent study that reported that the Wuhan-Hu-1-like virus can infect SD rats (16), although an earlier study showed that the prototype Wuhan-Hu-1-like SARS-CoV-2 cannot infect SD rats (6). Such a discrepancy may be due to variation in additional mutations in the challenge Wuhan-Hu-1-like strains or genetic variations in the SD rats used in these studies. Thus, further surveillance is needed to understand the virological prevalence in NYC rats, particularly for several emerging variants with high infectivity among rats, including those that circulated in NYC during the past two years of the COVID-19 pandemic.

A number of studies suggested that fragments of SARS-CoV-2 genomes were identified in sewage water systems, and that the prevalence of SARS-CoV-2 in sewage water systems coincides with outbreaks in resident human populations (17). However, no evidence has shown that SARS-CoV-2 viruses in sewage water is infectious (18), suggesting that sewage rats may have been exposed to the virus through unknown fomites, e.g. those contaminated with human food wastes. In a recent study, Zeiss et al. (13) showed that, in a controlled laboratory setting studying transmission of another rat respiratory beta coronavirus, SDAV, approximately one-quarter of naïve rats shed virus following fomite exposure. Notably, previously exposed seropositive rats became reinfected with SDAV at similar rates following fomite exposure 114-165 days later, indicating that immunity is temporary. Two of four rats (Rat #2 and #19) in our study were both seropositive and viral RNA-positive, implying that previously exposed seropositive animals may still contract and shed SARS-CoV-2, consistent with lack of sterilizing immunity in humans exposed to SARS-CoV-2 or rats given SDAV. These data imply that rats previously exposed to SARS-CoV-2 can still contribute to propagation of subsequent variants.

By using animal models, we further demonstrated that, in addition to Alpha and Beta variants reported earlier (6–9), Delta and Omicron variants can also cause robust infections in SD rats. The tested variants caused robust replication in both upper and lower respiratory tracts of rats, although they did not cause any body weight loss or other clinical signs. Of the three testing variants, Delta replicated the most efficiently. The omicron variant showed a lower viral replication than both Alpha and Delta, although the difference did not reach a statistically significant level between Omicron and Alpha. This finding is in line with earlier reports that Omicron replicated less efficiently and caused less lung pathology in wildtype or human ACE2 transgenic mice or hamsters compared with other variants (19, 20).

Structural modeling showed that all three variants Alpha, Delta and Omicron have enhanced binding to the rat ACE2 compared to the prototype Wuhan-Hu-1-like virus. In light of the biochemical data that Alpha and Delta RBDs bind to rat ACE2 equally well (14), the difference in the replication efficiency of the three viruses could be due to factors other than receptor binding affinity. It is also interesting to note that many RBD mutations observed in the three variants, such as N501Y in Alpha and L452R/T478K in Delta, interact with ACE2 residues that vary between human and rat/mouse (Fig. 4E). Therefore, rats and mouse likely play an important role in the evolution of Alpha, Delta, Omicron variants, as previously proposed by Zhang et al (15).

In addition to receptor binding, a number of other studies suggested that other structural and non-structural proteins may play a critical role in the viral replication *in vivo* and the host tropism of SARS-CoV-2 viruses. Syed et al. showed that, despite envelope protein substitutions inhibiting virus assembly, Omicron has an overall higher assembly efficiency than the original SARS-CoV-2, similar to Delta variant (21). Bojkova et al. showed that the Omicron variant is less effective in antagonizing the interferon response and has higher sensitivity in interferon treatment than the Delta variant which may be relevant with the substitutions on NSP3, NSP12, NSP13, nucleocapsid, and ORF3 proteins (22, 23). Of interest, Omicron did not have any observed amino acid substitutions throughout the course of virus challenge in SD rats, whereas Alpha and Delta did for spike, nucleoprotein, or non-structural proteins NSP6 and NSP13. The roles of these amino acid substitutions on virus fitness needs to be further studied.

In summary, we found that the rats in NYC sewage system have been exposed to SARS-CoV-2, and that the Delta and Omicron variants can infect rats in addition to the Alpha and Beta variants. Our findings highlight the potential risk of secondary zoonotic transmission from rats and the need for further monitoring of SARS-CoV-2 in wild rat populations.

## Materials and Methods

### Cells

Vero E6 cells (CRL-1586, American Type Culture Collection [ATCC]) and 293FT/hACE2+TMPRSS (17) were cultured in Dulbecco’s Modified Eagle medium (DMEM, Gibco) supplemented with 10% fetal bovine serum (FBS) at 37°C with 5% CO2. Rat lung epithelial cells L2 (CCl-149, ATCC) were cultured in F-12K Medium (ATCC) supplemented with 10% FBS at 37°C with 5% CO2. Rat primary tracheal epithelial cells (Cell Biologics) were grown on culture flasks or plates pre-coated with gelatin-based coating solution (Cell Biologics) in Complete Epithelial Cell Medium (Cell Biologics) at 37°C with 5% CO2.

### Viruses

The SARS-CoV-2/USA/CA_CDC_5574/2020 (B.1.1.7, NR-54011, BEI resources), and SARS-CoV-2/human/USA/MD-HP05285/2021 (B.1.617.2, NR-55671, BEI resources) were propagated on Vero E6 cells. The SARS-CoV-2/USA/MO-CV40709/2022 (BA.5.5, GIAID access No. EPI_ISL_15823386) were recovered from human nasopharyngeal swabs and propagated on Vero E6 cells.

### Virus challenge in rats

Six-week-old female SAS outbred Sprague Dawley (SD) rats (Charles River Laboratories) were housed in individually ventilated cages. SD rats were anesthetized with isoflurane, followed by intranasal inoculation with 2 × 10^4^ PFU/rat of SARS-CoV-2 diluted in 50 μl PBS. Clinical evaluation was performed daily, and body weight was determined daily through 10 dpi. At 2, 4, and 21 dpi, animals were euthanized for blood and tissue collection for seroconversion evaluation, viral load titration, and histology staining, respectively.

### Wild rat capture and sample collection

In the fall of 2021, APHIS Wildlife Services conducted sampling of Norway rats (*Rattus norvegicus*) in New York City (NYC) to look for evidence of SARS-CoV-2 infection. Methodology included trapping and collecting biological samples from rats around wastewater systems. Two trapping efforts during September and November were conducted with permission from the NYC Department of Parks and Recreation. Each effort consisted of three days of pre-baiting followed by four nights of trapping. Most animals were captured in city parks within the borough of Brooklyn, although some were captured near buildings outside of park boundaries. Once the animals were euthanized, biologists collected and processed fresh blood samples. Over the course of eight trapping nights, 79 rats were trapped and sampled. Blood samples along with the carcasses were shipped to the Wildlife Services National Wildlife Disease Program in Fort Collins, Co where tissues were extracted and sent to the University of Missouri for additional testing.

### Infectious viral titration by TCID_50_

Animal tissue were homogenized in DMEM with 0.3% bovine serum albumin (Sigma-Aldrich) and 1% penicillin/streptomycin (Gibco, Thermo Fisher Scientific) for 1 min at 6,000 rpm by using a homogenizer (Bertin, Precellys), and debris were pelleted by centrifugation for 10 min at 12,000 × *g*. Their infectious virus titers were determined by TCID_50_ with Vero E6 cells.

### Viral RNA detection

The RNA was extracted from the tissue homogenates by using GeneJet viral DNA/RNA purification kit (Thermo Fisher) or MagMax Pathogen RNA/DNA Kit (Thermo Fisher). The viral RNA was detected and quantified by qRT-PCR following the SARS-CoV-2 diagnosis panels by N1 (Forward primer sequence: 5’-GAC CCC AAA ATC AGC GAA AT-3’, Reverse primer sequence: 5’-TCT GGT TAC TGC CAG TTG AAT CTG-3’, Probe sequence: 5’-ACC CCG CAT TAC GTT TGG TGG ACC-3’) and/or N2 primer/probe mix (Forward primer sequence: 5’-TTACAAACATTGGCCGCAAA-3’, Reverse primer sequence: 5’-GCGCGACATTCCGAAGAA-3’, Probe sequence: 5’-ACAATTTGCCCCCAGCGCTTCAG-3’). The RT-qPCR was performed according to the manufacturer’s protocol using TaqMan Fast Virus 1-Step Master Mix (Thermo Fisher). Fluorescent signals were acquired using QuantStudio 6 Real-time PCR system (Thermo Fisher).

### Measurement of cytokine/chemokine expression

Total RNA was extracted from rat tissues by using a combination method of Trizol (Thermo Fisher Scientific) and RNeasy Mini kit (Qiagen) (24). The genomic DNA was removed by on-column DNase I (Qiagen) treatment during the RNA extraction. The RNA then was reverse transcribed into cDNA with SuperScript III Reverse Transcriptase (Thermo Fisher Scientific) with random hexamer primers (Thermo Fisher Scientific). The cDNA was used in real-time PCR with PowerUp SYBR Green Master Mix (Thermo Fisher Scientific) for specific targets (Table S4). The expression of housekeeping gene GAPDH was used to normalize the amount of RNA isolated from tissues. The 2^-ΔΔct^ methods were used to compare the differential gene expressions between testing samples. The mean fold change (2^-ΔΔCt^) values of triplicates and standard deviation are represented.

### Genome sequencing

SARS-CoV-2 whole-genome sequencing was performed by using QIAseq DIRECT SARS-CoV-2 Kit (QIAGEN). The quality of paired-end reads obtained from MiSeq sequencing was analyzed by using Qiagen CLC Genomics Workbench 22.0.1 and the Identify ARTIC V3 SARS-CoV-2 Low Frequency and Shared Variants (Illumina) workflow was used in genetic variant analyses. Nucleotide sequences were aligned using MAFFT v7.471, and the mutations were analyzed using nextclade (https://clades.nextstrain.org). Pan-viral target hybridization enrichment sequencing was performed by using RNA Prep with Enrichment (L) Tagmentation Kit (Illumina) and Comprehensive Viral Research Panel (Twist Biosciences).

### Phylogenetic analyses and molecular characterization

Time-scaled phylogenetic trees were generated using the two rat samples containing > 10% coverage (Rat# 2 and 19), five reference sequences for each variant of concern (Alpha, Beta, Delta, Gamma, and Omicron), five reference sequences for lineage A and lineage B viruses, and three randomly selected NYC sequences from each month. Phylogenetic analyses were performed using BEAST v2.7.0 with the Hasegawa, Kishino, and Yano (HKY)+⌈_4_ substitution model, an exponential coalescent growth prior, and a strict molecular clock. Independent runs were performed with chain lengths of 10,000,000 steps and sampled every 5,000 steps per run with a 10% burn-in. The resulting trees were summarized with TreeAnnotator and visualized using FigTree. A posterior probabilities cutoff of 0.70 was used to assess tree topology.

All publicly available sequences and associated metadata used in this dataset are published in GISAID’s EpiCoV database. All sequences in this dataset are compared relative to hCoV-19/Wuhan/WIV04/2019 (WIV04), the official reference sequence employed by GISAID (EPI_ISL_402124). Learn more at https://gisaid.org/WIV04. To view the contributors of each individual sequence with details such as accession number, virus name, collection date, originating lab and submitting lab, and the list of authors, please visit the doi listed with each dataset:

#### Data availability for GISAID samples included in our analyses

GISAID Identifier: EPI_SET_221019xq
doi: 10.55876/gis8.221019xq EPI_SET_221019xq is composed of 49 individual genome sequences. The collection dates range from 2019-12-24 to 2021-11-17; Data were collected in 11 countries and territories.

### Virus isolation

Virus isolation was done on Vero E6 cells, 293FT/hACE2+TMPRSS, L2, or rat primary tracheal epithelial cells. 200 μl of supernatant from homogenized tissues were mixed with an equal volume of cell culture medium and then inoculated onto pre-seeded cells in 6-well plates. After 1 hour of adsorption, the inoculum was removed, and the cells were washed with PBS and covered with fresh cell culture medium. The cells were monitored daily for cytopathogenic effects (CPE) and the supernatants were harvested at 3~5 dpi. The supernatants were inoculated to fresh cells for a maximum of 3 times until CPE was observed. The supernatants from the last inoculation were subjected to viral RNA extraction and SARS-CoV-2 specific real-time RT-PCR using SARS-CoV-2 diagnosis panels.

### ELISA

Anti-SARS-CoV-2 spike and anti-SARS-CoV-2 receptor binding domain (RBD) IgG or IgM antibodies were determined by using stabilized spike protein (NR-53524, BEI resources) or RBD (NR-53366, BEI resources) of SARS-CoV-2, respectively. The proteins were coated to 96-well ELISA plates (Nunc-Immuno, Thermo Scientific) at a concentration of 1 μg/ml in PBS. The plates were then blocked with 100 μl of 1% Bovine Serum Albumin (BSA, Research Products International) buffered in PBS containing 0.1% Tween 20 (PBST) and incubated at room temperature for 1 h. 1:100 diluted rat serum samples were added to the plates for 1 h incubation at 37 °C. After extensive washing with PBST, horseradish peroxidase (HRP)-conjugated goat anti-rat IgG (1:8,000, Thermo Scientific) or anti-rat IgM (1:8,000, Thermo Scientific) was added for 1 h at 37 °C. Following five-time washes with PBST, 100 μL of TMB-ELISA substrate (1-Step; Thermo Fisher Scientific) was added into each well. After 15 min incubation, the reaction was stopped by adding 100 μL of 1 M H_2_SO_4_ solution and optical densities were read at 450 nm (OD450) using Cytation 5 Cell Imaging Multimode Reader (Bio-Tek Instruments). Cutoff value was determined based on the mean background reactivity of all serum samples from naïve SD rats multiplied by 3.

### Microneutralization assay

The serum samples were heat-inactivated at 56 °C for 1 hour and then were two-fold serially diluted with a starting dilution of 1:5. The serum dilutions were mixed with equal volumes of 100 TCID_50_ of SARS-CoV-2 as indicated. After 1 h of incubation at 37 °C, 3.5 × 10^4^ Vero E6 cells were added into the serum-virus mixture in 96-well plates. The plates were incubated for 2 days at 37 °C in 5% CO_2_ and then the cells were fixed in 10% paraformaldehyde, penetrated by 0.1% TritonX-100, and strained with monoclonal rabbit antibody against SARS-CoV-2 nucleocapsid (Sino Biological). This was subsequently detected by the addition of HRP-conjugated goat anti-rabbit IgG (Thermo Fisher Scientific) and TMb-ELISA substrate (Thermo Fisher Scientific). OD_450_ was measured by Cytation 5 (Bio-Tek). The serum neutralizing titer is the reciprocal of the highest dilution resulting in an infection reduction of >50%.

### Structure modeling

The tertiary structure of the rat ACE2 (NP_001012006.1) was predicted by Alphafold2 using the Google colab server (https://colab.research.google.com/) (25). The RBD structure of alpha (SARS-CoV-2/USA/CA_CDC_5574/2020, B.1.1.7), delta (SARS-CoV-2/human/USA/MD-HP05285/2021, B.1.617.2) and omicron (SARS-CoV-2/USA/MO-CV40709/2022, BA.5.5) was taken from the PDB ID 7FBK, 7URQ, and 7XWA, respectively. To model the rat ACE2:RBD complex structure, rat ACE2 structural model and the structure of each of the three RDB domains were superposed onto their respective homologs in PDB ID 7XO9, the SARS-CoV-2 Omicron BA.2 variant RBD complexed with human ACE2 (26) using Pymol (The PyMOL Molecular Graphics System, Version 2.0 Schrödinger, LLC). The resulting complex structures were subjected to energy minimization using Phenix (27). Structure figures were prepared using Pymol.

### Statistical analysis

Statistical significance was tested using a one-way ANOVA with Tukey’s multiple comparisons by Graphpad Prism 9.1.0.

### Ethics statement

Rats were captured in Brooklyn under a wildlife damage management agreement between USDA/APHIS Wildlife Services and the New York City Department of Parks and Recreation. The animal experiments were performed under the protocol number #38742 approved by the Care and Use of Laboratory Animals of the University of Missouri per the USDA Animal Welfare Regulations. All experiments involved with live viruses were performed in an approved biosafety level 3 (BSL-3) or animal biosafety level 3 (ABSL-3) facility at the Laboratory of Infectious Diseases, University of Missouri-Columbia under protocol #20-14 in compliance with the Institutional Biosafety Committee of the University of Missouri-Columbia.

## Acknowledgments

This study was supported by USDA American Rescue Plan funding. The authors thank George Sarafianos, Rebecca Patterson, Haley Hudson for their assistance in this study. We thank Marc Johnson for suggesting the wastewater systems targeted in this study and Mark Jacking, John Pistone, Raven Shuman, Deana Brabant Oatman, Maxwell Tanner, Jack Ramirez, Allen Gosser, Bobby Corrigan, Tim Linder and Tom Gidlewski for for wild rat capture, sample collection and necropsy/tissue processing. We also thank Samantha Gerb, Sarah Schlink, Charles Moley, and Shakera Fudge for their technical supports in animal experiments.

The findings and conclusions in this publication are those of the author(s) and should not be construed to represent any official USDA or U.S. Government determination or policy.

## References

1. Medicine JHUo. 2022. Coronavirus Resource Center. https://coronavirus.jhu.edu/data. Accessed March 4.

2. Cui S, Liu Y, Zhao J, Peng X, Lu G, Shi W, Pan Y, Zhang D, Yang P, Wang Q. 2022. An Updated Review on SARS-CoV-2 Infection in Animals. Viruses 14.

3. Chandler JC, Bevins SN, Ellis JW, Linder TJ, Tell RM, Jenkins-Moore M, Root JJ, Lenoch JB, Robbe-Austerman S, DeLiberto TJ, Gidlewski T, Kim Torchetti M, Shriner SA. 2021. SARS-CoV-2 exposure in wild white-tailed deer (Odocoileus virginianus). Proc Natl Acad Sci U S A 118.

4. Hale VL, Dennis PM, McBride DS, Nolting JM, Madden C, Huey D, Ehrlich M, Grieser J, Winston J, Lombardi D, Gibson S, Saif L, Killian ML, Lantz K, Tell RM, Torchetti M, Robbe-Austerman S, Nelson MI, Faith SA, Bowman AS. 2022. SARS-CoV-2 infection in free-ranging white-tailed deer. Nature 602:481–486.

5. Hammer AS, Quaade ML, Rasmussen TB, Fonager J, Rasmussen M, Mundbjerg K, Lohse L, Strandbygaard B, Jorgensen CS, Alfaro-Nunez A, Rosenstierne MW, Boklund A, Halasa T, Fomsgaard A, Belsham GJ, Botner A. 2021. SARS-CoV-2 Transmission between Mink (Neovison vison) and Humans, Denmark. Emerg Infect Dis 27:547–551.

6. Shuai H, Chan JF, Yuen TT, Yoon C, Hu JC, Wen L, Hu B, Yang D, Wang Y, Hou Y, Huang X, Chai Y, Chan CC, Poon VK, Lu L, Zhang RQ, Chan WM, Ip JD, Chu AW, Hu YF, Cai JP, Chan KH, Zhou J, Sridhar S, Zhang BZ, Yuan S, Zhang AJ, Huang JD, To KK, Yuen KY, Chu H. 2021. Emerging SARS-CoV-2 variants expand species tropism to murines. EBioMedicine 73:103643.

7. Pan T, Chen R, He X, Yuan Y, Deng X, Li R, Yan H, Yan S, Liu J, Zhang Y, Zhang X, Yu F, Zhou M, Ke C, Ma X, Zhang H. 2021. Infection of wild-type mice by SARS-CoV-2 B. 1.351 variant indicates a possible novel cross-species transmission route. Signal Transduct Target Ther 6:420.

8. Zhang C, Cui H, Li E, Guo Z, Wang T, Yan F, Liu L, Li Y, Chen D, Meng K, Li N, Qin C, Liu J, Gao Y, Zhang C. 2022. The SARS-CoV-2 B.1.351 Variant Can Transmit in Rats But Not in Mice. Front Immunol 13:869809.

9. Montagutelli X, Prot M, Levillayer L, Salazar EB, Jouvion G, Conquet L, Beretta M, Donati F, Albert M, Gambaro FJB. 2021. Variants with the N501Y mutation extend SARS-CoV-2 host range to mice, with contact transmission. BioRxiv.

10. Bosco-Lauth AM, Root JJ, Porter SM, Walker AE, Guilbert L, Hawvermale D, Pepper A, Maison RM, Hartwig AE, Gordy P, Bielefeldt-Ohmann H, Bowen RA. Survey of peridomestic mammal susceptibility to SARS-CoV-2 infection. doi:https://doi.org/10.1101/2021.01.21.427629.

11. Miot EF, Worthington BM, Ng KH, de Lataillade LG, Pierce MP, Liao Y, Ko R, Shum MH, Cheung WY, Holmes EC, Leung KS, Zhu H, Poon LL, Peiris MJ, Guan Y, Leung GM, Wu JT, Lam TT. 2022. Surveillance of Rodent Pests for SARS-CoV-2 and Other Coronaviruses, Hong Kong. Emerg Infect Dis 28:467–470.

12. Colombo VC, Sluydts V, Marien J, Vanden Broecke B, Van Houtte N, Leirs W, Jacobs L, Iserbyt A, Hubert M, Heyndrickx L, Goris H, Delputte P, De Roeck N, Elst J, Arien KK, Leirs H, Gryseels S. 2022. SARS-CoV-2 surveillance in Norway rats (Rattus norvegicus) from Antwerp sewer system, Belgium. Transbound Emerg Dis 69:3016–3021.

13. Zeiss CJ, Asher JL, Vander Wyk B, Allore HG, Compton SR. 2021. Modeling SARS-CoV-2 propagation using rat coronavirus-associated shedding and transmission. PLoS One 16:e0260038.

14. Yao W, Ma D, Wang H, Tang X, Du C, Pan H, Li C, Lin H, Farzan M, Zhao J, Li Y, Zhong G. 2021. Effect of SARS-CoV-2 spike mutations on animal ACE2 usage and in vitro neutralization sensitivity. https://www.biorxivorg/content/101101/20210127428353v3fullpdf.

15. Zhang W, Shi K, Geng Q, Ye G, Aihara H, Li F. 2022. Structural basis for mouse receptor recognition by SARS-CoV-2 omicron variant. Proc Natl Acad Sci U S A 119:e2206509119.

16. Yu D, Long Y, Xu L, Han JB, Xi J, Xu J, Yang LX, Feng XL, Zou QC, Qu W, Lin J, Li MH, Yao YG. 2022. Infectivity of SARS-CoV-2 and protection against reinfection in rats. Zool Res 43:945–948.

17. Smyth DS, Trujillo M, Gregory DA, Cheung K, Gao A, Graham M, Guan Y, Guldenpfennig C, Hoxie I, Kannoly S, Kubota N, Lyddon TD, Markman M, Rushford C, San KM, Sompanya G, Spagnolo F, Suarez R, Teixeiro E, Daniels M, Johnson MC, Dennehy JJ. 2022. Tracking cryptic SARS-CoV-2 lineages detected in NYC wastewater. Nat Commun 13:635.

18. Robinson CA, Hsieh HY, Hsu SY, Wang Y, Salcedo BT, Belenchia A, Klutts J, Zemmer S, Reynolds M, Semkiw E, Foley T, Wan X, Wieberg CG, Wenzel J, Lin CH, Johnson MC. 2022. Defining biological and biophysical properties of SARS-CoV-2 genetic material in wastewater. Sci Total Environ 807:150786.

19. Halfmann PJ, Iida S, Iwatsuki-Horimoto K, Maemura T, Kiso M, Scheaffer SM, Darling TL, Joshi A, Loeber S, Singh G, Foster SL, Ying B, Case JB, Chong Z, Whitener B, Moliva J, Floyd K, Ujie M, Nakajima N, Ito M, Wright R, Uraki R, Warang P, Gagne M, Li R, Sakai-Tagawa Y, Liu Y, Larson D, Osorio JE, Hernandez-Ortiz JP, Henry AR, Ciuoderis K, Florek KR, Patel M, Odle A, Wong LR, Bateman AC, Wang Z, Edara VV, Chong Z, Franks J, Jeevan T, Fabrizio T, DeBeauchamp J, Kercher L, Seiler P, Gonzalez-Reiche AS, Sordillo EM, Chang LA, van Bakel H, et al. 2022. SARS-CoV-2 Omicron virus causes attenuated disease in mice and hamsters. Nature 603:687–692.

20. Yuan S, Ye ZW, Liang R, Tang K, Zhang AJ, Lu G, Ong CP, Man Poon VK, Chan CC, Mok BW, Qin Z, Xie Y, Chu AW, Chan WM, Ip JD, Sun H, Tsang JO, Yuen TT, Chik KK, Chan CC, Cai JP, Luo C, Lu L, Yip CC, Chu H, To KK, Chen H, Jin DY, Yuen KY, Chan JF. 2022. Pathogenicity, transmissibility, and fitness of SARS-CoV-2 Omicron in Syrian hamsters. Science 377:428–433.

21. Syed AM, Ciling A, Taha TY, Chen IP, Khalid MM, Sreekumar B, Chen PY, Kumar GR, Suryawanshi R, Silva I, Milbes B, Kojima N, Hess V, Shacreaw M, Lopez L, Brobeck M, Turner F, Spraggon L, Tabata T, Ott M, Doudna JA. 2022. Omicron mutations enhance infectivity and reduce antibody neutralization of SARS-CoV-2 virus-like particles. Proc Natl Acad Sci U S A 119:e2200592119.

22. Bojkova D, Widera M, Ciesek S, Wass MN, Michaelis M, Cinatl J, Jr. 2022. Reduced interferon antagonism but similar drug sensitivity in Omicron variant compared to Delta variant of SARS-CoV-2 isolates. Cell Res 32:319–321.

23. Bojkova D, Rothenburger T, Ciesek S, Wass MN, Michaelis M, Cinatl J, Jr. 2022. SARS- CoV-2 Omicron variant virus isolates are highly sensitive to interferon treatment. Cell Discov 8:42.

24. Untergasser A. 2008. RNAprep - Trizol combined with Columns. http://www.molbi.de/protocols/rna_prep_comb_trizol_v1_0.htm. Accessed April 13.

25. Mirdita M, Schutze K, Moriwaki Y, Heo L, Ovchinnikov S, Steinegger M. 2022. ColabFold: making protein folding accessible to all. Nat Methods 19:679–682.

26. Xu Y, Wu C, Cao X, Gu C, Liu H, Jiang M, Wang X, Yuan Q, Wu K, Liu J, Wang D, He X, Wang X, Deng SJ, Xu HE, Yin W. 2022. Structural and biochemical mechanism for increased infectivity and immune evasion of Omicron BA.2 variant compared to BA.1 and their possible mouse origins. Cell Res 32:609–620.

27. Liebschner D, Afonine PV, Baker ML, Bunkoczi G, Chen VB, Croll TI, Hintze B, Hung LW, Jain S, McCoy AJ, Moriarty NW, Oeffner RD, Poon BK, Prisant MG, Read RJ, Richardson JS, Richardson DC, Sammito MD, Sobolev OV, Stockwell DH, Terwilliger TC, Urzhumtsev AG, Videau LL, Williams CJ, Adams PD. 2019. Macromolecular structure determination using X-rays, neutrons and electrons: recent developments in Phenix. Acta Crystallogr D Struct Biol 75:861–877.

