## Supplementary material for "SARS-CoV-2 exposure in Norway rats (*Rattus norvegicus*) from New York City": Table S1-S4

[1]Center for Influenza and Emerging Infectious Diseases, University of Missouri, Columbia, Missouri, USA; [2]Department of Molecular Microbiology and Immunology, School of Medicine, University of Missouri, Columbia, Missouri, USA; [3]Bond Life Sciences Center, University of Missouri, Columbia, Missouri, USA; [4]USDA APHIS Wildlife Services National Wildlife Disease Program, Fort Collins, Colorado, USA; [5]USDA APHIS Wildlife Services, Fort Collins, Colorado, USA; [6]Viral Diseases Branch, Walter Reed Army Institute of Research, Silver Spring, Maryland, USA; [7]Department of BioSciences, Rice University, Houston, TX, USA; [8]School of Medicine, Yale University, New Haven, Connecticut, USA; [9]Department of Electrical Engineering & Computer Science, College of Engineering, University of Missouri, Columbia, Missouri, USA.

* Xiu-Feng Wan; Thomas J. DeLiberto

**Table S1.** SARS-CoV-2 whole genome sequencing.

| Rat code | Number of Reads | Reads mapped to SARS-CoV-2 reference (%) | SARS-CoV-2 Genome Coverage (%) | Average Depth of Coverage |
|---|---|---|---|---|
| #2 | 7,826,926 | 92.97 | 21.3 | 28,924.70 |
| #19 | 10,423,292 | 93.42 | 18.4 | 38,495.70 |
| #43 | 100,528 | 75.78 | 1.9 | 376.10 |
| #46 | 1,811,062 | 98.11 | 1.6 | 6,274.80 |

**Table S2.** Coronavirus sequences identified using pan-viral target hybridization enrichment sequencing.

| Rat code | Number of sequence reads | Coronavirus detected | Virus reads | Identified genes |
|---|---|---|---|---|
| 2 | 1582668 | SARS-CoV-2 | 13 | nsp 2, 3,10,14, S, ORF3a, E |
| 19 | 1328920 | SARS-CoV-2 | 10 | nsp 1, 3, 5, 13, 14, ORF6, ORF7a |
| 43 | 2747254 | SARS-CoV-2 | 18 | nsp 2-5, 10, 12, 13, 15, N |
| 30 | 1137027 | Rat coronavirus | 97 | nsp1-5, 7, 8, 12, 13, 15, 16, S, M, N, 3’UTR |
| 38 | 1026796 | SARS-CoV-2 | 24 | nsp 2-4, 9-16, S |
| 46 | n.a. | n.a. | n.a. | n.a. |

**Table S3.** Amino acid substitutions among the rats challenged with SARS-CoV-2 variants.

| Animal ID | Amino acid residue on viral proteins[b] | | | | | | |
|---|---|---|---|---|---|---|---|
| | ORF1a | ORF1b | Spike | | | | N |
| | 3718[c] | 979[d] | 19 | 74 | 681 | 950 | 63 |
| Alpha | | | | | | | |
| Challenge strain | V | D | T | N | P | D | D |
| SD10 (2 dpi)[a] | V | **N** | T | **K** | P | D | D |
| SD11 (2 dpi) | V | **N** | T | **K** | P | D | D |
| SD12 (2 dpi) | V | D | T | **K** | P | D | D |
| SD13 (4 dpi) | V | **N** | T | **K** | P | D | D |
| SD14 (4 dpi) | V | **N** | T | **K** | P | D | D |
| SD15 (4 dpi) | V | **N** | T | **K** | P | D | D |
| Delta | | | | | | | |
| Challenge strain | V | D | R | N | P | D | D |
| SD19 (2 dpi) | **A** | D | R | N | **R** | **N** | D |
| SD20 (2 dpi) | **A** | D | R | N | **R** | **N** | D |
| SD21 (2 dpi) | **A** | D | R | N | **R** | **N** | D |
| SD22 (4 dpi) | **A** | D | **T** | N | **R** | **N** | D |
| SD23 (4 dpi) | **A** | D | **T** | N | **R** | **N** | **G** |
| SD24 (4 dpi) | **A** | D | **T** | N | **R** | **N** | **G** |
| Omicron | | | | | | | |
| Challenge strain | V | D | T | N | P | D | D |
| SD28 (2 dpi) | V | D | T | N | P | D | D |
| SD29 (2 dpi) | V | D | T | N | P | D | D |
| SD30 (2 dpi) | V | D | T | N | P | D | D |
| SD31 (4 dpi) | V | D | T | N | P | D | D |
| SD33 (4 dpi) | V | D | T | N | P | D | D |

Note: [a]dpi, days post infection; [b]the amino acid substitutions compared to the challenge strain were highlighted in red; [c] residue 148 in NSP6, proofreading 3′-5′ exonuclease, N7-methyltransferase; [d] residue 56 in NSP13, replication organelle formation.

**Table S4.** Primers used to quantify mRNA expression of proinflammatory markers

| Gene | Forward Primer (5′-3′) | Reverse Primer (5′-3′) |
|---|---|---|
| GAPDH | GAGACAGCCGCATCTTCTTG | TGACTGTGCCGTTGAACTTG |
| IL-1α | TCGGGAGGAGACGACTCTAA | GAAAGCTGCGGATGTGAAGT |
| IL-1β | CTGTGACTCGTGGGATGATG | GGGATTTTGTCGTTGCTTGT |
| IFN-γ | GGAACTGGCAAAAGGACGGTA | CTCGAACTTGGCGATGCTCA |
| IFN-α | CTGGTGGTGATGAGCTACTGG | TTGTGCCAGGAGTGTGAAGG |
| IFN-β | AACCTCAGCTACAGGACGGA | TGGAGCATCACTTGAATGGCA |
| IL-6 | CTCATTCTGTCTCGAGCCCA | CTGTGAAGTCTCCTCTCCGG |
| IL-10 | GCAGGACTTTAAGGGTTACTTGG | GGGGAGAAATCGATGACAGC |
| TNF-α | CAAACCACCAAGCAGAGGAG | GAGGCTGACTTTCTCCTGGT |
| CCL-2 | AGCCAACTCTCACTGAAGCC | TGGGGCATTAACTGCATCTGG |
| IP-10 | CCGCATGTTGAGATCATTGCC | AGACCTTCTTTGGCTCACCG |
